# Effects of bromodomain protein inhibition in a mouse model of Niemann-Pick type C disease

**DOI:** 10.64898/2026.06.24.734200

**Authors:** Martina Parente, Amélie Barthelemy, Sara Caputo, Noémie Charlery-Adèle, Christian Galmiche, Claudia Tonini, Simona Lobasso, Danilo Prtvar, Sabina Tahirovic, Sophie Reibel Foisset, Valentina Pallottini, Frank W. Pfrieger

**Affiliations:** Department of Science, Section Biomedical Science and Technology, University Roma Tre, Viale Marconi 446, 00146 Rome, Italy; Centre National de la Recherche Scientifique, Université de Strasbourg, Institut des Neurosciences Cellulaires et Intégratives, 8 allée du Général Rouvillois, 67000 Strasbourg, France; Chronobiotron, CNRS, UAR 3415, 67000 Strasbourg, France; Department of Translational Biomedicine and Neuroscience, University of Bari Aldo Moro, piazza G. Cesare 11, 70124 Bari, Italy; German Center for Neurodegenerative Diseases (DZNE) Munich, 81377 Munich, Germany; Neuroendocrinology, Metabolism and Neuropharmacology Unit, LIFE Institute for Research and Health Care - Santa Lucia IRCCS, Via del Fosso Fiorano 64, 00143 Rome, Italy

**Keywords:** Cholesterol/animal model, cholesterol/trafficking, cholesterol/metabolism, drug therapy, epigenetic regulation, lysosome, rare disease, proteostasis, bromodomain, BET protein inhibition, histone acetylation

## Abstract

Defects in lysosomal lipid handling provoke fatal disorders presenting neurovisceral symptoms with variable onset and life spans. A prime example is Niemann-Pick type C disease (NPCD), where export of cholesterol and other lipids from the endosomal-lysosomal system is impaired due to variants of either NPC intracellular cholesterol transporter 1 (NPC1) or NPC intracellular cholesterol transporter 2 (NPC2). Therapeutic options for NPCD are limited to palliative care and disease-modifying drugs, and there is an unmet need for new treatments. Based on positive effects in patient-derived fibroblasts *in vitro*, we explored here how inhibition of bromodomain and extra-terminal domain (BET) proteins, a family of histone acetylation readers, affects a well-established mouse model bearing the frequent I1061T variant of NPC1. To this end, we used employed intraperitoneal administration of the blood-brain-barrier penetrating drug JQ1 or vehicle (dimethylsulfoxide), and gauged drug effects based on weight, biomarker levels and quantitative assessment of motor coordination. Treatment of mice with JQ1 induced beneficial effects on some outcome measures, notably a transient increase in weight, reduction of liver cholesterol, and improvements in the footprint test, but these effects occurred only in male but not in female mice. Our results indicate bromodomain proteins as potential therapeutic drug target for NPCD and reveal sex-dependent BET protein signaling in mice.

## Background

Niemann-Pick type C disease (NPCD) is a rare, pan-ethnic, autosomal-recessive lysosomal disorder presenting progressive and ultimately fatal neurovisceral symptoms [1–4]. The primary cause are variants of *NPC intracellular cholesterol transporter 1* (*NPC1*; OMIM #<u>257220</u>; 95% of cases) [5, 6] or of *NPC intracellular cholesterol transporter 2* (*NPC2*; OMIM <u>607625</u>; 5% of cases) [7]. Other genetic and epigenetic factors modify disease progression and severity [8–10]. The ubiquitously expressed proteins reside in the membrane (NPC1) [11, 12] and lumen (NPC2) of late endosomes [7, 13, 14]. Dysfunction of either protein causes intracellular accumulation of unesterified cholesterol [15, 16] and impairs lysosomal [17–20] and mitochondrial function [21–25]. The mechanisms causing dysfunction and degeneration of specific cell types, notably in the liver, lung and brain, are not fully understood [26].

At present, there is an unmet need for therapeutic approaches. Current options are limited to disease-modifying treatment with N-butyl-deoxynojirimycin (OGT918, Miglustat, Zavesca) [27], arimoclomol (Miplyffa) in combination with Miglustat [28] or N-acetyl-L-leucine (Levacetylleucine) [29, 30]. Here, we explored inhibition of bromodomain and extra-terminal (BET) proteins as a new therapeutic approach for NPCD using a well-established knock-in mouse line expressing a frequent pathogenic variant of NPC1 (NPC1^tm(I1061T)Dso^) [31] that is degraded due to misfolding [32–34]. Homozygous mutant mice present progressive weight loss and neurovisceral symptoms starting at 8 weeks of age and premature death between 14 to 17 weeks of age [31, 35–40]. Our therapeutic targets, BET proteins, bind to acetylated lysines on histones and thereby regulate gene transcription [41]. The translational potential of their inhibition is supported by positive effects in cardiometabolic disease [42] and different types of cancer [43–48] observed in recent clinical trials and studies on human tissues. Previously, we showed that JQ1, a well-established competitive inhibitor of BET proteins [49], raises the level of NPC1 protein in cultured cells [50, 51]. This effect has a potential therapeutic effect for NPC1 variants with residual function. In skin fibroblasts from NPCD patients, JQ1 diminished lysosomal expansion and cholesterol accumulation, and augmented extracellular release of lysosomal components in a dose-, time- and patient-dependent manner [51]. These results prompted us to launch the next step of preclinical testing, to explore beneficial effects of JQ1 in an established NPCD mouse model in vivo.

## Methods

### Animals and drug administration

A colony of transgenic mice (B6.129-*Npc1*^tm1Dso^/J) was maintained by crossing heterozygous animals (Chronobiotron, UAR 3415, Strasbourg, France). Genotypes of animals were determined by PCR from tail biopsies using established primers (ProtocolID 28838 for strain #027704; Jackson Laboratory). Groups of 5-to 6-weeks-old homozygous mutant and wildtype littermates were transferred to the animal facility in Rome, housed in Makrolon cages (26.7 cm x 20.7 cm x 14 cm), under controlled conditions (temperature 20–21 °C, 55–65% relative humidity and a 12/12h light cycle with lights on at 7 am). No inclusion or exclusion criteria were established, animals were assigned arbitrarily to treatment groups, and no experimenter blinding was applied as objective physical measures were taken. JQ1 was dissolved in dimethyl sulfoxide (DMSO) for injection volumes of 500 µl/kg body weight and administered daily on 6 days per week by intraperitoneal injections for 48 hours or 4 weeks at doses of 20 or 40 mg/kg as indicated. Control animals were treated with vehicle (DMSO). Treatment duration of four weeks was a compromise between cost and time required for progressive manifestation of neurologic symptoms and possible drug effects. All behavioral experiments were performed between 10 am to 12 pm/noon. All animals were sacrificed at 10 am on Wednesdays to allow for two prior daily drug administrations. Except for few initial experiments, the sex of individual animals was taken into account as key biological determinant [52, 53]. The experiments were approved by the Italian Ministry of Health (Rome, Italy; Authorization N° 677/2020-PR) and performed in agreement with the ARRIVE (Animals in Research: Reporting In Vivo Experiments) guidelines, the guidelines of the Italian Ministry of Health (D.L. 26/14) and the European Community Directive 2010/63/EU.

### Western blots

Tissues from different organs and brain areas were lysed in homogenization buffer [sucrose 0.1 M, KCl 0.05 M, KH_2_PO_4_ 0.04 M, EDTA 0.04 M, pH 7.4, with 1:1,000 protease inhibitor cocktail (#P8340) and 1:400 phosphatase inhibitor cocktail (#P0044); Sigma-Aldrich/Merck]. Spleen, liver and brain samples were prepared at 1:10 and 1:5 w/v respectively, to yield total lysates. Samples were homogenized using a Potter homogenizer followed by sonication (VCX 130 PB, Sonics, Newtown, 06470 CT) on ice for 20s. Samples were then centrifuged at 13,000 rpm for 10 min at 4 °C to remove cell debris. Protein concentration was determined using the Bradford assay (Bio-Rad Laboratories, Hercules, CA, USA). For immunoblotting, samples were diluted with 4x sample buffer (20% SDS, Tris HCl 1 M pH 6.8, 0.01% Bromophenol Blu 0.01%, 100% β-mercaptoethanol and 100% glycerol), heated at 100 °C for 5 minutes, loaded on 7-10% acrylamide gel and subjected to SDS-PAGE (40 µg of protein/lane). Proteins were transferred to nitrocellulose membranes (Trans-Blot Turbo Transfer System; Bio-Rad Laboratories, Milan, Italy). Membranes were blocked with fat-free milk or bovine serum albumin (BSA) (5% in Tris-buffered saline 0.138M NaCl, 0.027M KCl, 0.025M Tris-HCl, and 0.05% Tween-20, pH 7.6) for 1h at room temperature (RT), and probed at 4°C overnight with the following primary antibodies: NPC1 (Novus Biologicals, NB400-148; dilution 1:2500), cMYC (Santa Cruz, sc-40; 1:500), SREBP2 (Santa Cruz, sc13552; 1:1000), CYP3A4 (Abcam, ab3572; 1:2000), calbindin (Antibodies.com, A85360; 1:1000), and vinculin (Sigma-Aldrich, V9131; 1:40.000) or beta-tubulin (Sigma-Aldrich, T9026; 1:40.000) as loading controls. Subsequently, membranes were incubated for 1h with horseradish peroxidase-conjugated secondary IgG antibodies (Bio-Rad Laboratories, Milan, Italy). Protein-antibody immunocomplexes on nitrocellulose filters were reacted with Clarity Western ECL substrate (Bio-Rad Laboratories, Milan, Italy, #1705061). Chemiluminescence signals were acquired with the ChemiDoc MP system (Bio-Rad Laboratories, Milan, Italy) and images were analyzed by ImageJ. Uncut images of Western blots are provided in Supplementary Fig. 1.

### Other molecular analyses

For liver cholesterol, livers were dissected and stored at −80 °C. Concentrations of free and total cholesterol were determined using a cholesterol assay kit following the manufacturer’s instructions (#CS0005, Sigma-Aldrich/Merck). For neurofilament light chain (NFL), up to 0.5 mL of trunk blood were collected per animal into 2 mL plastic tubes, and spun down (10 min; 3000 g at 4°C) within a maximum of 30 min from collection. Subsequently, the supernatant was collected and stored at −80°C until further use. The serum concentration of NFL was measured using the NF-light Advantage Assay Kit (Quanterix) and the Simoa HD-X Analyser (Quanterix) following the manufacturer’s instructions.

### Behavioral tests

Several tests were explored to assess motor coordination in mice from our colony. Animals were subjected to these tests at 8 weeks of age, before receiving drug or vehicle, and at 12 weeks of age, after drug or vehicle administration.

### Footprint test

To obtain footprints, the hindfeet of mice were coated with black, non-toxic paint, and the animals were allowed to walk along a 40-cm-long, 6-cm-wide runway leading to an enclosed box. All mice underwent two training runs followed by three test runs. For each run, a new sheet of white paper was placed on the floor of the runway. Footprint patterns were characterized by four quantitative gait parameters: (1) step distance of right and left hind paws defined as the average distance of forward movement between consecutive paw placements, (2) stride distance defined as the distance between the two hindfeet, (3) the number of steps taken within a 20 cm segment of the runway measured for each of the three test runs, and (4) the base of support defined as the lateral distance between the two hindfeet during walking. For each measure, a minimum of three values was obtained per run, excluding footprints at the beginning and end of the run, where the animal was initiating and terminating movement. At least three runs were performed per animal. Unless indicated otherwise, mean values across the three runs were used for subsequent analysis.

### Balance beam test

Mice were trained to cross a 5 mm wide and 80 cm long beam with square profile suspended at 50 cm height. For tests, mice were allowed to run three times across the beam. Three parameters were obtained: the average time to cross the entire beam, the running length, and the number of slips while crossing the beam. For animals that did not cross the beam entirely, the length was noted and the time was set at 20s.

### Rotarod

For the five-station rotarod performance test (Mouse Rotarod, Ugo Basile, Italy), mice were placed on a 0.03 m wide rod rotating horizontally at constant speed (5 rounds per minute) that was positioned low enough (0.16 m) to prevent injuries, but high enough to permit discernible falls. The time until animals fell down (latency) was recorded. A maximum latency of 60s was defined for mice that did not fall. Adaptation runs were performed during the week before the experiments and 1-minute-long resting intervals were inserted between runs.

### Data analysis and visualisation

Data analysis and visualisation were accomplished using ImageJ, the open source software R (version 4.5.1.) [54] and selected R packages [coin v. 1.4.3. [55]; data.table v. 1.17.8 [56], DHARMa v. 0.5.0 [57], estimated marginal means (emmeans) v 1.11.2.8.) [58], ggplot2 v. 4.0.0 [59], glmmTMB [60], lme4 v. 1.1.37 [61], lmerTest v. 3.1.3 [62], PCAtest v. 0.0.2 [63]]. Datasets with sample sizes >3 are shown by boxplots, for n = 3, datasets are shown by mean and standard deviation. In all figures, symbols represent values of individual animals. In Western blots, only single factor effects (sex, genotype or drug) on two experimental groups could be compared for technical reasons, and the Wilcoxon-Mann-Whitney test was used for statistical analysis. PCA was performed using the prcomp command of base R with prior centering and scaling of data. Principal components were compared by two-way analysis of variance (ANOVA) of base R. To analyse impact of sex, genotype and drug treatment as categorical fixed factors on weight and behavioral outcome measures, we used various general linear, linear mixed and generalized linear models with variable-dependent distributions (see Supplementary Data 1). For mixed models, animal set (cohorts) and individual unique animal identifiers were included as random factors. To validate the models with respect to zero-inflation, heteroscedasticity, uniformity and dispersion of residuals, we used the DHARMa package. To calculate estimated marginal means based on the fitted models and to perform selected pairwise post-hoc tests we used the emmeans library. Original data (Supplementary Data 1) and details of statistical analyses (Supplementary Data 2) are provided in accompanying files.

## Results

To reveal effects of BET protein inhibition *in vivo*, we used homozygous transgenic knock-in mice NPC1^tm(I1061T)Dso^, further referred to as NP or mutant mice, and their wildtype (WT) littermates. To inhibit BET proteins, we administered JQ1, a prototype inhibitor that passes the blood-brain barrier [64], by daily intraperitoneal injections. Control animals run in parallel received vehicle (DMSO) injections.

### In vivo exploration of JQ1 effects upon short-term treatment of NPC1 I1061T mutant mice

First, we tested whether short-term treatment of mice with JQ1 affects the levels of NPC1, cMYC [65] and SREBP2 [66] that are known targets of BET protein regulation. To this end, 8-weeks-old mutant and wildtype mice received daily intraperitoneal injections of JQ1 at 20 mg/kg or vehicle (DMSO). After two days of treatment, we determined protein levels in cerebellum, cortex, liver, and spleen by immunoblotting. Short-term treatment with JQ1 affected the levels of the three proteins with tissue- and protein-dependent manner. This includes an increase and a decrease of NPC1 in the cerebellum and liver of mutant mice, respectively, and selective increases of cMYC and SREBP2 in the spleen and cortex, respectively (Fig. 1; Supplementary Data 1, 2). Despite their divergence, the observed changes encouraged us to further explore how JQ1 affects mutant and wildtype mice in vivo.

**Figure 1.**
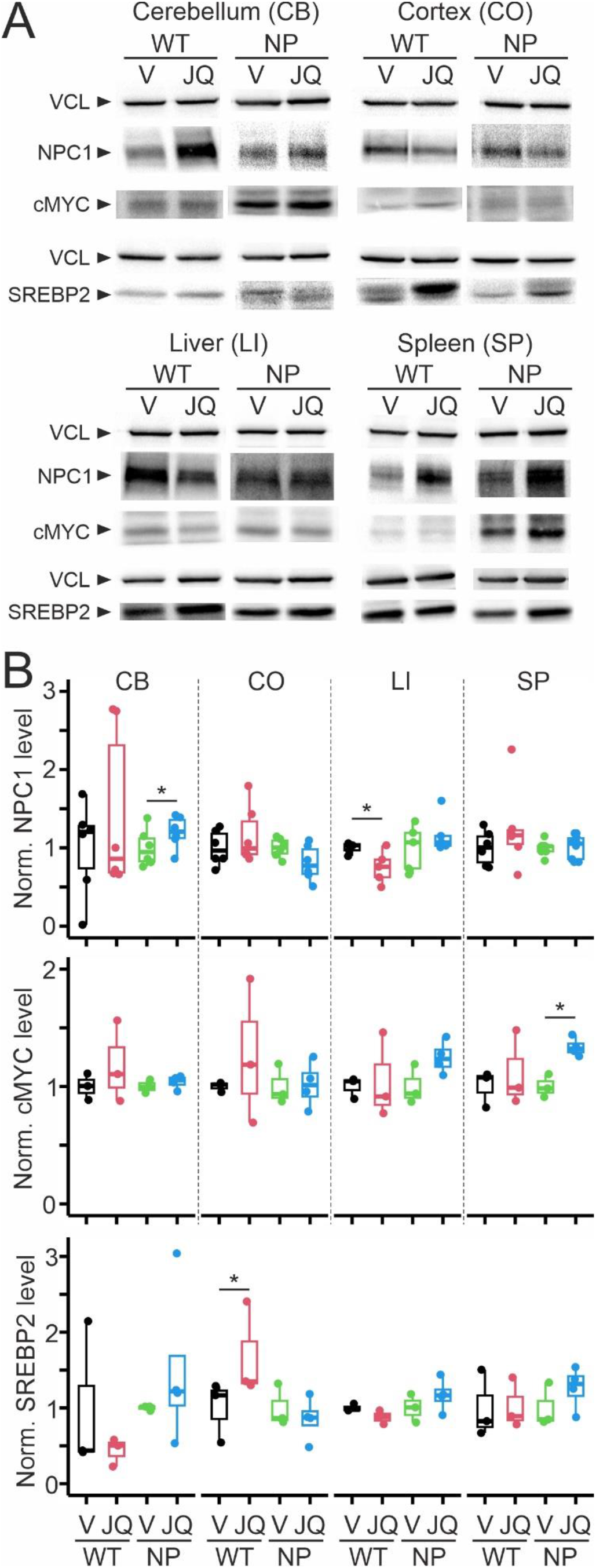
In vivo effects of short-term treatment with JQ1 on NPC1, cMYC and SREBP2 in selected organs and brain regions of wildtype and mutant mice. **A**, Representative Western blots showing levels of NPC1, cMYC and SREBP2 with vinculin (VCL) as loading control in indicated organs and brain regions of wildtype (WT) or mutant (NP) mice treated at 8 weeks for 48h using daily intraperitoneal injections with JQ1 (20 mg/kg) or DMSO (V, vehicle). B, Boxplots showing normalized levels of indicated proteins from indicated tissues of WT and NP mice treated with JQ1 or V. Values were normalized first to loading controls and then to levels of vehicle-treated animals per genotype analysed on the same blot. Asterisks indicate statistically significant differences between vehicle- and JQ1-treated animals for specific proteins and tissues (* p < 0.05; Wilcoxon-Mann-Whitney test; n = 3–7 animals per group; see Supplementary Data 1, 2 for data and analyses).

### Lower weights and impaired motor control in mutant mice at 8 weeks of age

Primary neurovisceral signs of NPC1 dysfunction in mice are progressive weight loss and decline of motor coordination [9, 31, 67–73]. We aimed to start treatment at neurologic disease onset to align with clinical reality, as most NPCD patients receive diagnosis and disease-modifying therapies after presentation of neurologic symptoms [4]. Previous studies assessed motor control in I1061T mice by diverse tests including gait analysis [35, 38], beam balance [37, 40], rotarod [31, 37, 38, 40], coat hanger and vertical screen [40]. They showed impaired motor control in this NPCD mouse model already at 8 weeks of age compared to wildtype controls [35, 37, 38].

To test whether mutant mice from our colony present neurologic symptoms at this age, we subjected naïve (untreated) mice to three independent behavioral tests, namely footprint, beam balance and rotarod providing multiple outcome measures per animal. We first analysed this rich multi-parametric dataset by principal component analysis (PCA) to determine whether weight and behavioral measures reveal changes that distinguish mutant and wildtype mice at 8 weeks of age. Evaluation by a random permutation test [63, 74] revealed significance of the first three principal components encouraging further exploration (Supplementary Data 1, 2). As shown in Fig. 2, the first principal component (PC1) covered 29.8% of the variance and clearly separated genotypes both visually and statistically without sex-specific differences. The component had strongest loadings from three measures of the footprint test (Fig. 2D). PC2 covered 20.2% of variance with strong loadings from body weight, footprint and beam balance measures showing significant differences between wildtype and mutant mice only in males. PC3 covered 15.4% of the variance with loadings from beam balance measures, but this component did not exhibit differences between genotypes (Fig. 2A–C).

**Figure 2.**
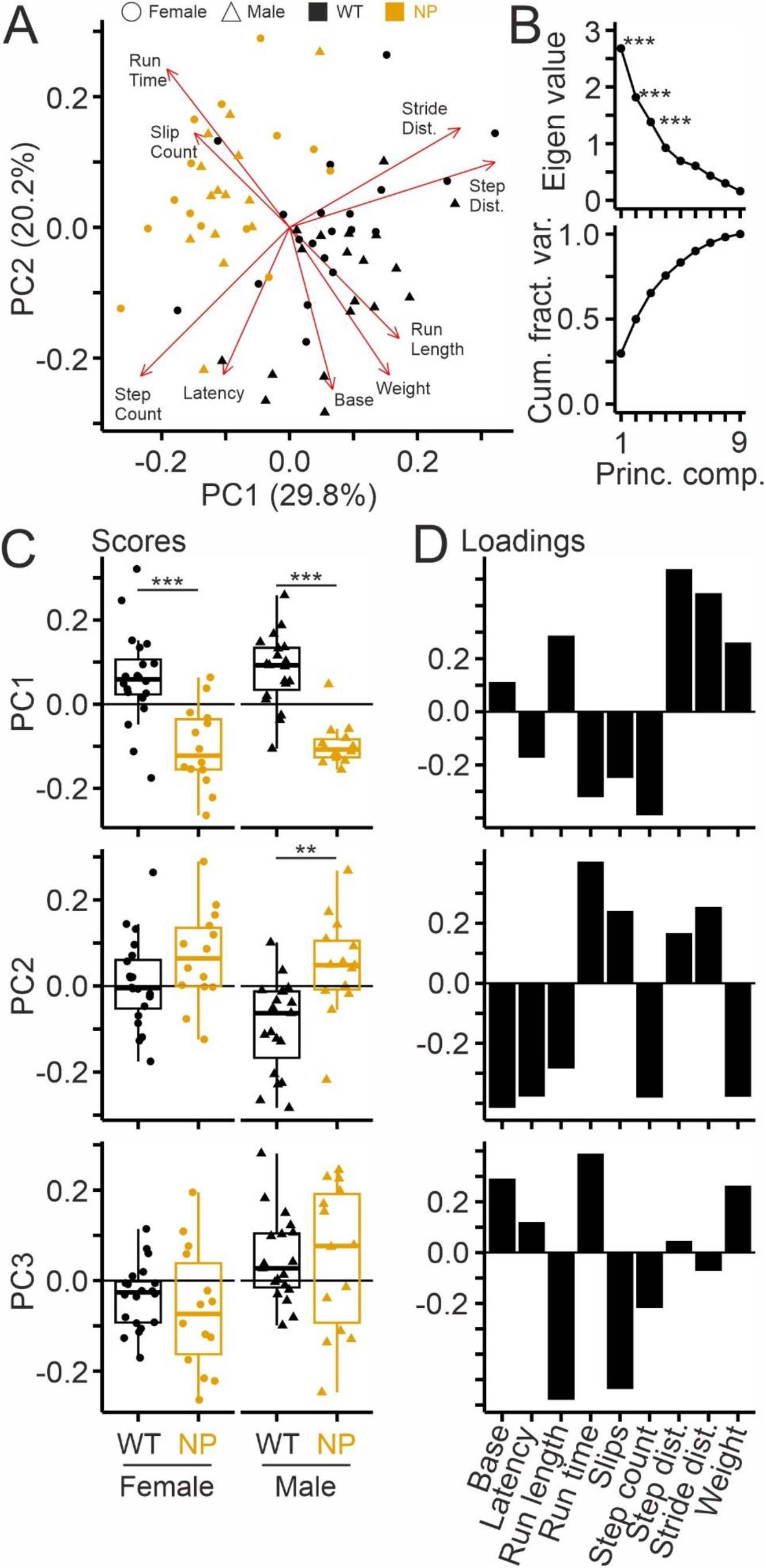
Principle component analysis revealing genotype- and sex-specific differences in mice at 8 weeks of age. A, Biplot showing scores (symbols) and loadings (arrows) of the top three principal components (PC) summarizing weight and indicated behavioral test measures from individual female (circles) and male (triangles) mutant (orange) and wildtype (black) mice subjected to footprint, beam balance and rotarod tests at 8 weeks of age. B, Screeplot (top) and line plot (bottom) showing Eigenvalues of PCs and cumulative relative variance explained by each PC, respectively. C, Boxplots showing scores of the indicated components in the indicated animal groups. D, Column plots showing loadings from each test measure to indicated principal components. Asterisks in B and D indicate significant PCs and loadings, respectively (* p < 0.05, ***, p < 0.001; 100 bootstrap replicates; 100 random permutations; PCAtest). Asterisks in C indicate significant differences between genotypes (*** p < 0.001; * p < 0.05; two-way ANOVA; Tukey’s pairwise posthoc test; female WT/NP: n = 20/14; male WT/NP: 19/14; see Supplementary Data 1, 2 for data and analyses).

**Suppl. Fig. 2.**
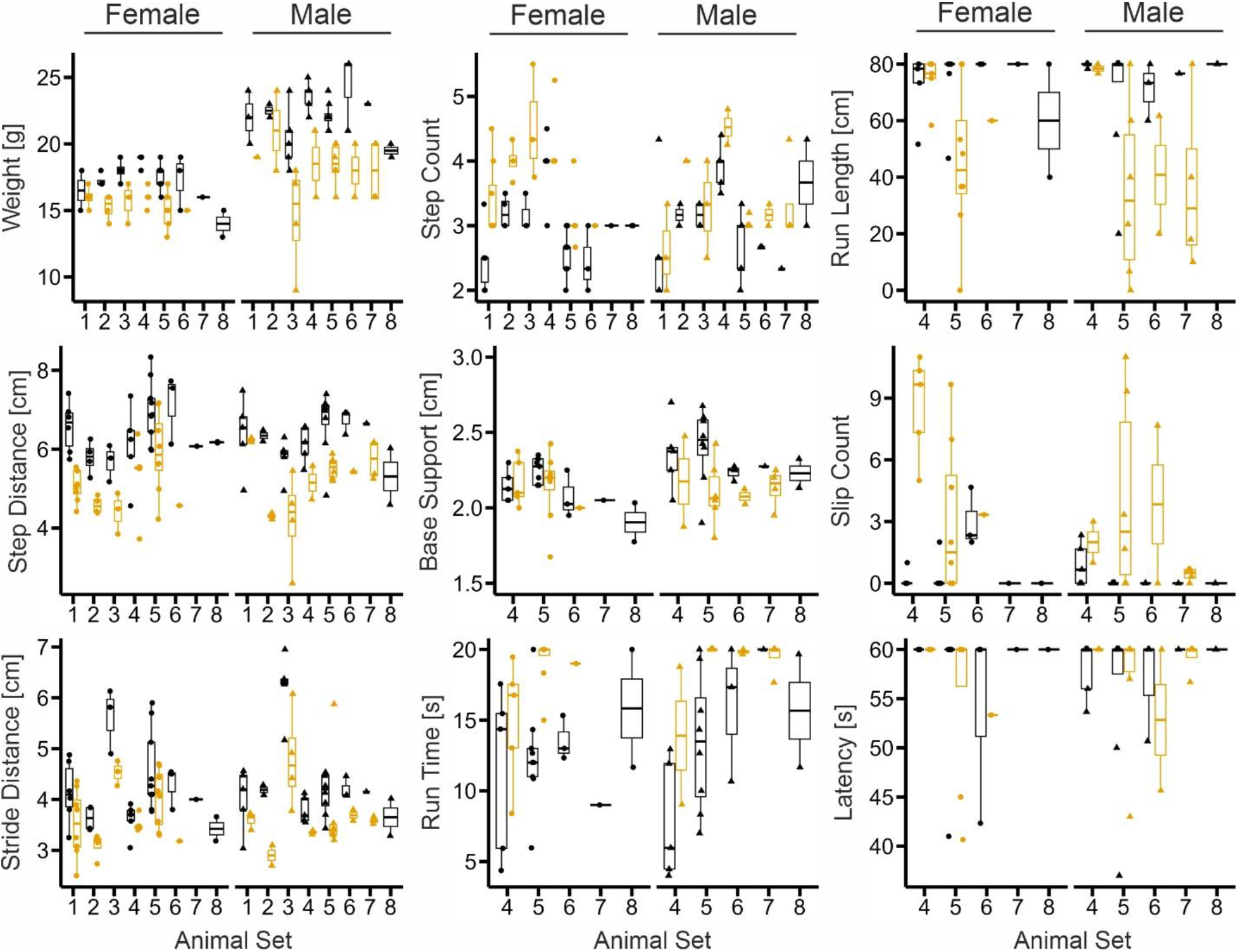
Weights and measures from behavioral tests in mutant and wildtype mice at 8 weeks of age. Boxplots showing weight and measures from footprint (step distance, stride distance, step count, base of support), beam balance (run time, run length, slip count) and rotarod test (latency) in female (circles) and male (triangles) mutant (orange) and wildtype (black) mice at 8 weeks of age separated by animal cohort. Note differences in some measure levels among cohorts. Symbols represent means from individual animals.

We next analysed outcome measures individually. As we noticed variation among the cohorts of animals that were tested sequentially (Supplementary Fig. 2), we explored sex- and genotype-dependent differences using generalized linear mixed models with variable-dependent response distributions. We included sex and genotype as fixed factors and animal cohort and – in some cases – unique animal identifiers as random variables (Supplementary Data 1, 2). Compared to their wildtype littermates, eight-weeks-old mutant mice had lower weights (Fig. 3) and scored worse in nearly all behavioral measures regardless of their sex. Exceptions were the base of support from the footprint test, which showed significant genotype x sex interaction, and the latency from the rotarod test, which showed no main effects. These results indicated that both footprint and beam balance test are sufficiently sensitive to detect motor defects in untreated male and female mice expressing the I1061T variant already at 8 weeks of age.

**Figure 3.**
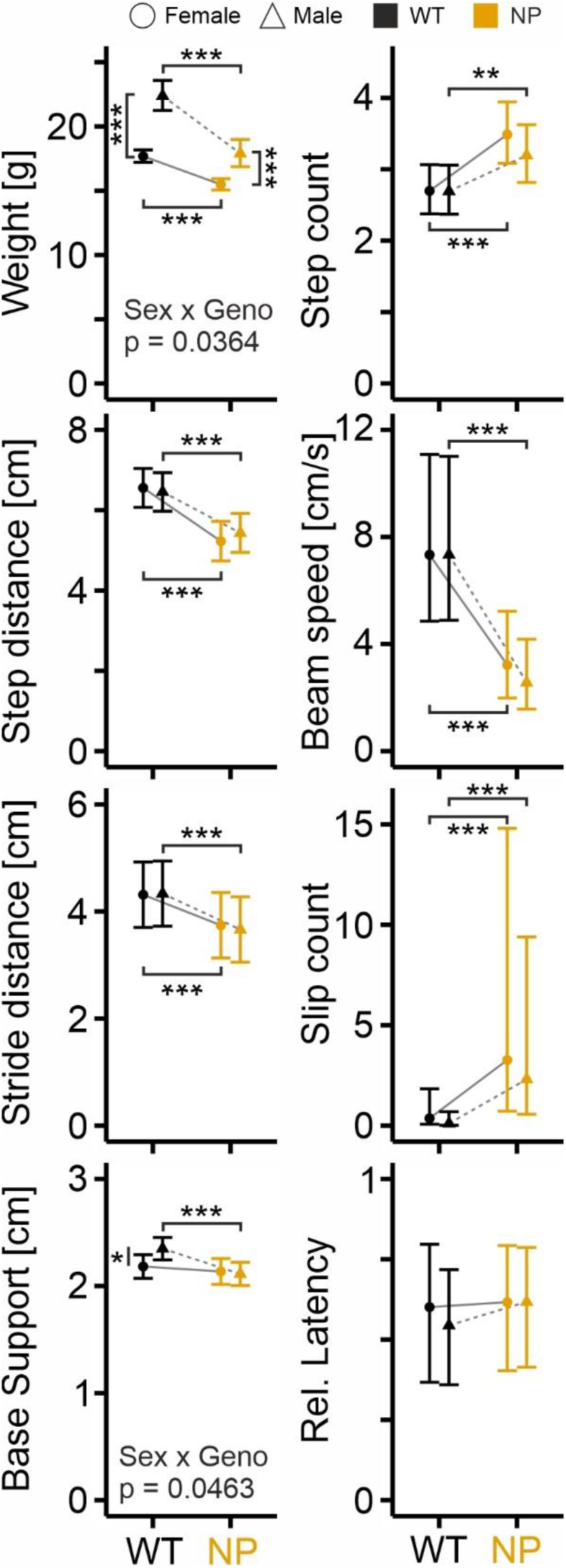
Genotype- and sex-dependent differences in weight and outcome measures of behavioral tests in mice at 8 weeks of age. Estimated marginal means of indicated measures analysed by measure-specific generalized linear mixed models (see Supplementary Data 1, 2 for data and analyses). Depending on the distribution, measures were back-transformed to the response scale. Whiskers indicate 95% confidence intervals. Asterisks designate statistically significant differences between indicated groups (*, p < 0.05; **, p < 0.01; ***, p < 0.001; pairwise post-hoc tests with Tukey’s adjustment. Sample sizes: weight: female WT/NP: n = 27/25; male WT/NP: 27/21; footprint measures: female WT/NP: 30/29; male WT/NP: 28–29/22–23; footprint base of support: female WT/NP: n = 17/14; male WT/NP: 17/14; beam balance and rotarod: WT/NP: 17/13–14; male WT/NP: 17/13–14).

**Suppl. Fig. 3.**
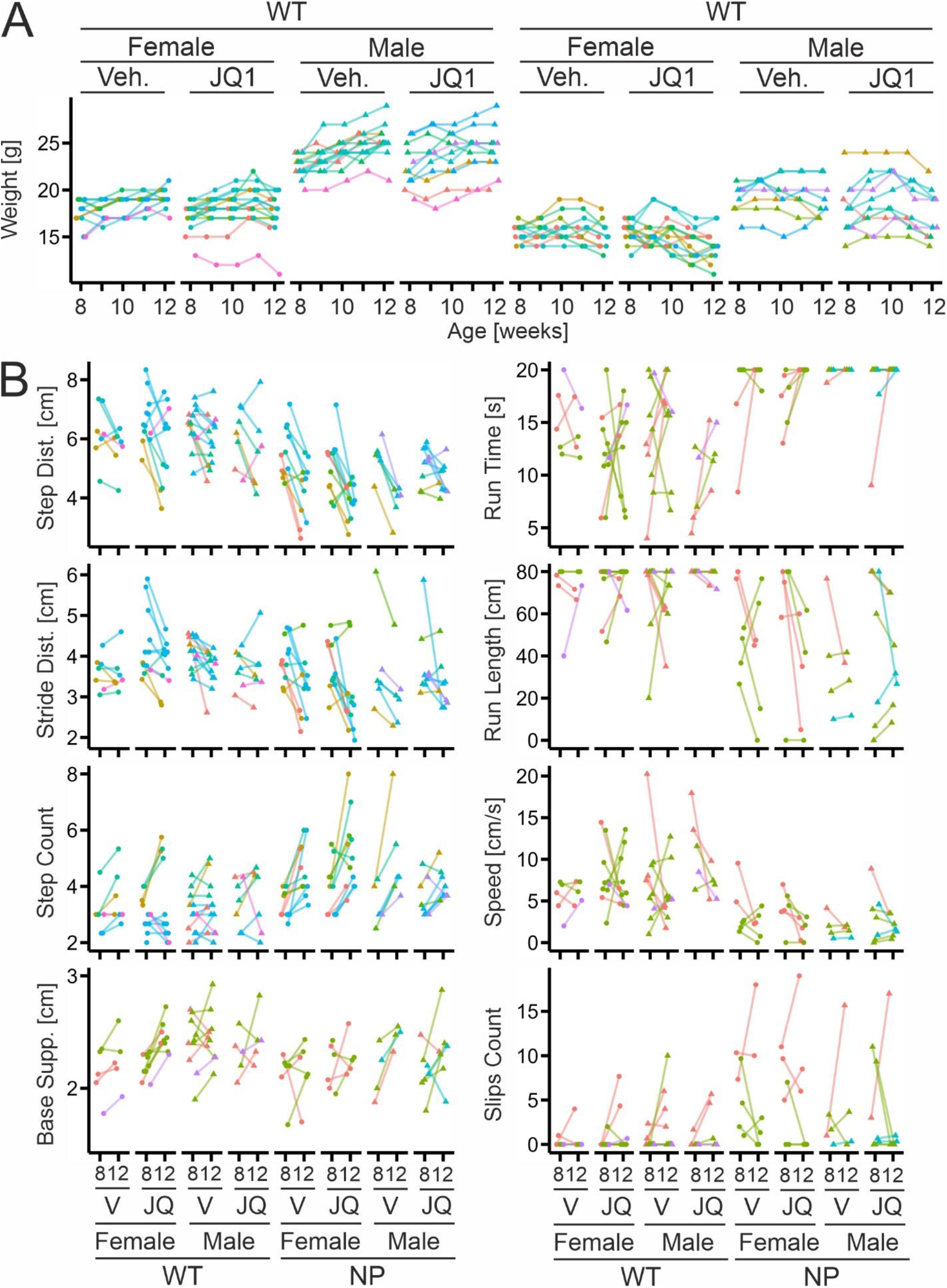
Weights and outcome measures of behavioral tests in mutant and wildtype mice treated with vehicle or drug. A, Line plots showing weights of individual animals during the 4-weeks-long treatment period separated by genotype, sex and treatment. Colors indicate different animal cohorts. B, Line plots showing measures from individual animals obtained by footprint (step distance, stride distance, step count, base of support), beam balance (run time, run length, speed, slip count) and rotarod test (latency) at 8 and 12 weeks of age separated by genotype, sex and treatment. Symbols represent mean values from each animal obtained from 3-5 test runs. Colors indicate different animal cohorts.

### Effects of JQ1 on body weight and motor control in mutant and wildtype mice after long-term treatment

To evaluate whether long-term treatment with JQ1 has positive effects on NPC1 mutant mice *in vivo*, we employed a before/after design (Fig. 4A). Starting at 8 weeks of age (before measure), we treated mutant animals and their wildtype littermates for four weeks with JQ1 or vehicle (DMSO) by daily intraperitoneal injections, and we monitored their body weight weekly, and behavioral tests were performed again at 12 weeks of age (after measure). Given the modest changes induced by short-term treatment (Fig. 1) we increased the dose to 40 mg/kg for long-term treatment. As observed for 8 weeks data, the weights differed individually and among animal cohorts (Supplementary Fig. 3A). We therefore analysed the repeated weight measures of WT and NP mice separately with a linear mixed model including treatment, age and sex as fixed factors and animal cohorts and unique animal identifiers as random factors (Supplementary Data 1, 2). In male mutant mice, JQ1 induced a weight gain until 10 weeks of age, which vanished during subsequent weeks (Fig. 4B). In female mutant mice, the drug induced an age-dependent weight loss (Fig. 4B). In wildtype animals, JQ1 did not affect the age-dependent weight gain of males, but it halted the weight gain in females during the last week of treatment leading to a significant treatment-induced weight difference at 12 weeks of age (Fig. 4B).

**Figure 4.**
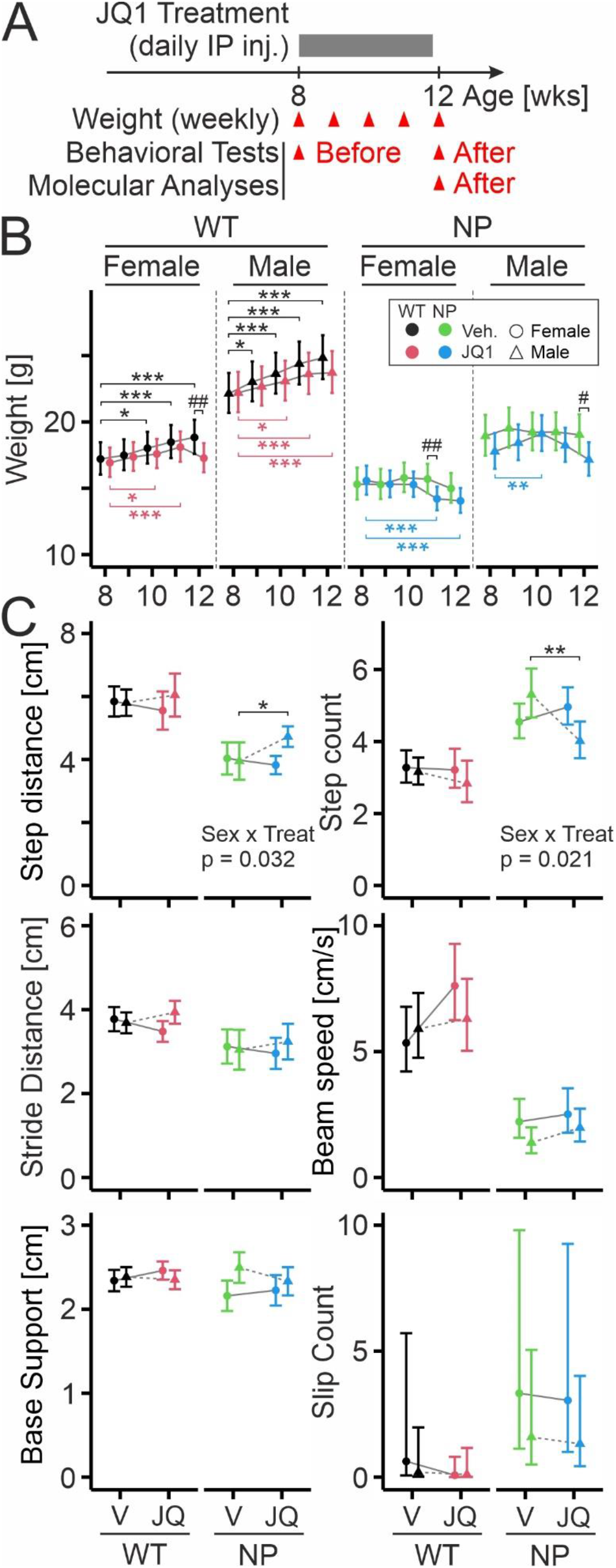
Effects of JQ1 on body weight and motor control in mutant and wildtype mice after long-term treatment. A, Diagram illustrating our experimental before/after design to test drug effects: animals were treated starting from 8 weeks of age for four weeks by daily intraperitoneal injections of JQ1 (40 mg/kg) or vehicle (DMSO). Body weights were measured weekly during the treatment period. Behavioral tests were performed at age 8 (before; see Fig. 2, 3) and 12 (after). Molecular analyses were performed at the end of the treatment period. B, Estimated marginal means (symbols) and 95% confidence intervals (whiskers) of body weights in animals separated by genotype and sex at indicated ages during treatment with vehicle or drug. Values were estimated by a generalized linear mixed model (see Supplementary Data 2) and back-transformed to the response scale. Asterisks indicate statistically significant changes compared to 8 weeks of age per sex and treatment (*, p < 0.05; **, p < 0.01; ***, p < 0.001). Number symbols indicate statistically significant changes between treatments per sex and age (#, p < 0.05; ##, p < 0.01; pairwise post-hoc tests with Tukey adjustment; sample sizes: WT: female Veh/JQ1: n = 11/16; male Veh/JQ1: 13/11; NP: female Veh/JQ1: 11/13; male Veh/JQ1: 8/10; see Supplementary data 1, 2 for data and analyses). C, Estimated marginal means (symbols) and 95% confidence intervals (whiskers) of indicated measures from footprint and beam balance tests in mice at 12 weeks of age with indicated sex and genotype after treatment with JQ1 or vehicle. Values were estimated by variable-dependent generalized linear (mixed) models (Supplementary Data 2) and back-transformed to the response scale. Asterisks indicate statistically significant differences (*, p < 0.05; **, p < 0.01; pairwise post-hoc tests with Tukey adjustment; sample sizes: footprint measures: WT: female Veh/JQ1 n = 10/14; male Veh/JQ1: 13/11; NP: female Veh/JQ1: 11/13; male Veh/JQ1: 8/10; beam balance measures: WT: female Veh/JQ1 n = 8/11; male Veh/JQ1: 10/9; NP: female Veh/JQ1: 5/6; male Veh/JQ1: 6/7).

We next assessed how JQ1 affects motor control using those measures from the footprint and beam balance test that showed significant genotype-specific differences in untreated animals at 8 weeks of age (Fig. 2, 3). For the beam balance test, we calculated the speed by dividing run distance by run time. As shown in Supplementary Fig. 3B, the measures showed considerable interindividual variability. To reveal drug effects, we used different general linear (mixed) models with sex and treatment as fixed factors and with variable-dependent response distributions and link functions (Supplementary Data 2). As shown in Fig. 4C, JQ1 improved performance in the footprint test in mutant mice as indicated by a significant increase in step distance and a decreased the number of steps compared to vehicle. Interestingly, these changes were seen only in male, but not in female mice (Fig. 4C). In wildtype animals, JQ1 did not show significant effects except for a spurious increase in the speed of females during the beam balance (Fig. 4C). Taken together, our results reveal some positive effects of JQ1 in male mutant mice including a transient weight gain as well as an increased step distance and concomitantly decreased step counts in the footprint test.

### Effects of JQ1 on molecular markers after long-term treatment

We next tested whether the positive JQ1-induced changes seen in male mutant mice were accompanied by changes of molecular markers. To this end, we measured levels of NPC1, cMYC and SREBP2 in different tissues of mutant mice. We separated males and females to test whether sex-specific drug effects on behavioral parameters were reflected by corresponding changes in the selected proteins (Fig. 5). Effects of JQ1 on NPC1 levels were variable, but the drug robustly decreased levels of the target protein cMYC across tissues in male animals. In female mice, the effect was weaker and did not reach statistical significance (Fig. 5A, B). Similarly, JQ1 also decreased levels of SREBP2 in liver and spleen compared to vehicle and this effect reached statistical significance in males only (Fig. 5C). These findings indicated coherent drug-induced changes in weight, footprint test and protein levels. The fact that we measured levels of all three proteins in liver lysates of individual animals allowed us to test whether levels of these three proteins were correlated (Fig. 5D). Interestingly, the levels of NPC1 and of cMYC were negatively correlated in the liver and positively in the spleen, whereas levels of SREBP2 and cMYC were positively correlated in both tissues (Figure 5D). This observation indicated tissue-specific effects of BET protein inhibition on target proteins.

**Figure 5.**
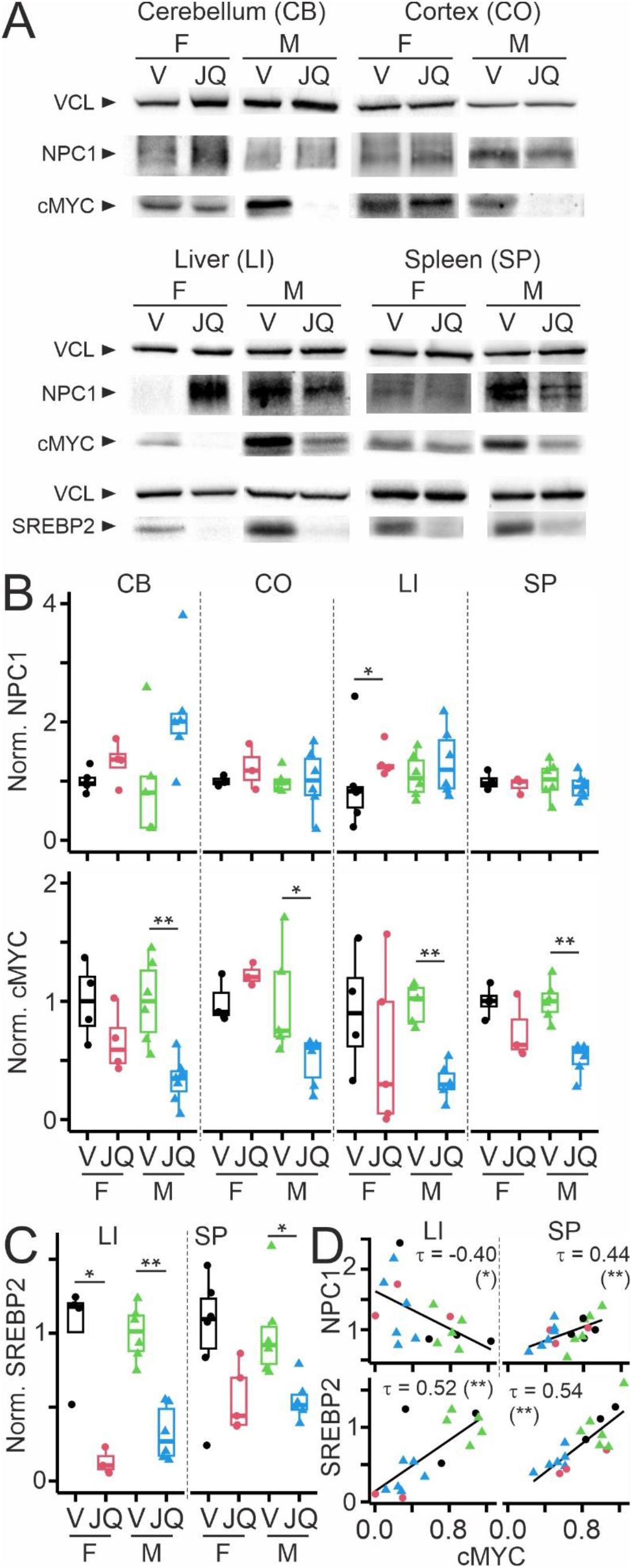
*In vivo* effects of long-term treatment with JQ1 on NPC1, cMYC and SREBP2 in selected organs and brain regions of mutant mice. A, Representative Western blots showing levels of NPC1, cMYC and SREBP2 with vinculin (VCL) as loading control in indicated brain regions and organs of female (F) and male (M) mutant mice treated at 8 weeks of age for 4 weeks with JQ1 (40 mg/kg) or DMSO (V, vehicle) by daily intraperitoneal injections. B, Boxplots showing normalized levels of indicated proteins from indicated tissues of female (F) and male (M) mutant mice treated with JQ1 or vehicle (V). Values were normalized first to loading controls and then to levels of vehicle-treated animals per genotype. C, Normalized levels of SREBP2 in liver (LI) and spleen (SP) from female (F) and male (M) mutant mice after treatment with JQ1 or vehicle. In B and C, asterisks indicate statistically significant differences (*, p < 0.05; **, p < 0.01; Wilcoxon-Mann-Whitney test; n = 3–7 animals per group). D, Scatterplots showing normalized protein levels of NPC1 (top) and SREBP2 (bottom) in liver (LI) and spleen (SP) plotted against corresponding cMYC levels in individual mutant mice as shown in panels B and C (same color and symbol codes). Numbers indicate correlation coefficients τ and asterisks indicate statistically significant correlations (Kendall rank correlation test; *, p < 0.05 and ** p < 0.01; NPC1 vs. cMYC: LI n = 18, SP n = 19 animals; SREBP2 vs. cMYC: LI/SP 16/18 animals).

We next measured other disease-relevant molecules. Cholesterol accumulation in the liver has been observed in I1061T mice [31] and other murine models of NPCD [67, 72, 73, 75–77]. JQ1 reduced the liver content of total and free cholesterol only in males, but not in female animals in agreement with weight, footprint test and protein data (Fig. 6A). With respect to markers reporting neurodegeneration in NPCD [78], protein levels of NFL in serum (Fig. 6B) and of calbindin in the cerebellum (Fig. 6C) were increased and decreased, respectively in untreated mutant mice compared to wildtype littermates as expected from previous studies [79]. Compared to vehicle, JQ1 showed a tendency to increase serum levels of NFL notably in female mutant mice (Fig. 6B) and to increase the level of calbindin in mutant males (Fig. 6C). Together, these findings provided further evidence for sex-specific effects of BET protein inhibition. To address a possible reason, we measured the levels of CYP3A in the liver. This enzyme metabolizes JQ1 [80] and shows strong expression in female mice [81]. Indeed, in 12-weeks-old untreated mice, female wildtype mice had higher levels of CYP3A compared to male littermates supporting a possible impact on drug availability. However, in mutant mice of the same age, this sex-specific difference was not present, and CYP3A levels seemed unaffected by JQ1 (Fig. 6D).

**Figure 6.**
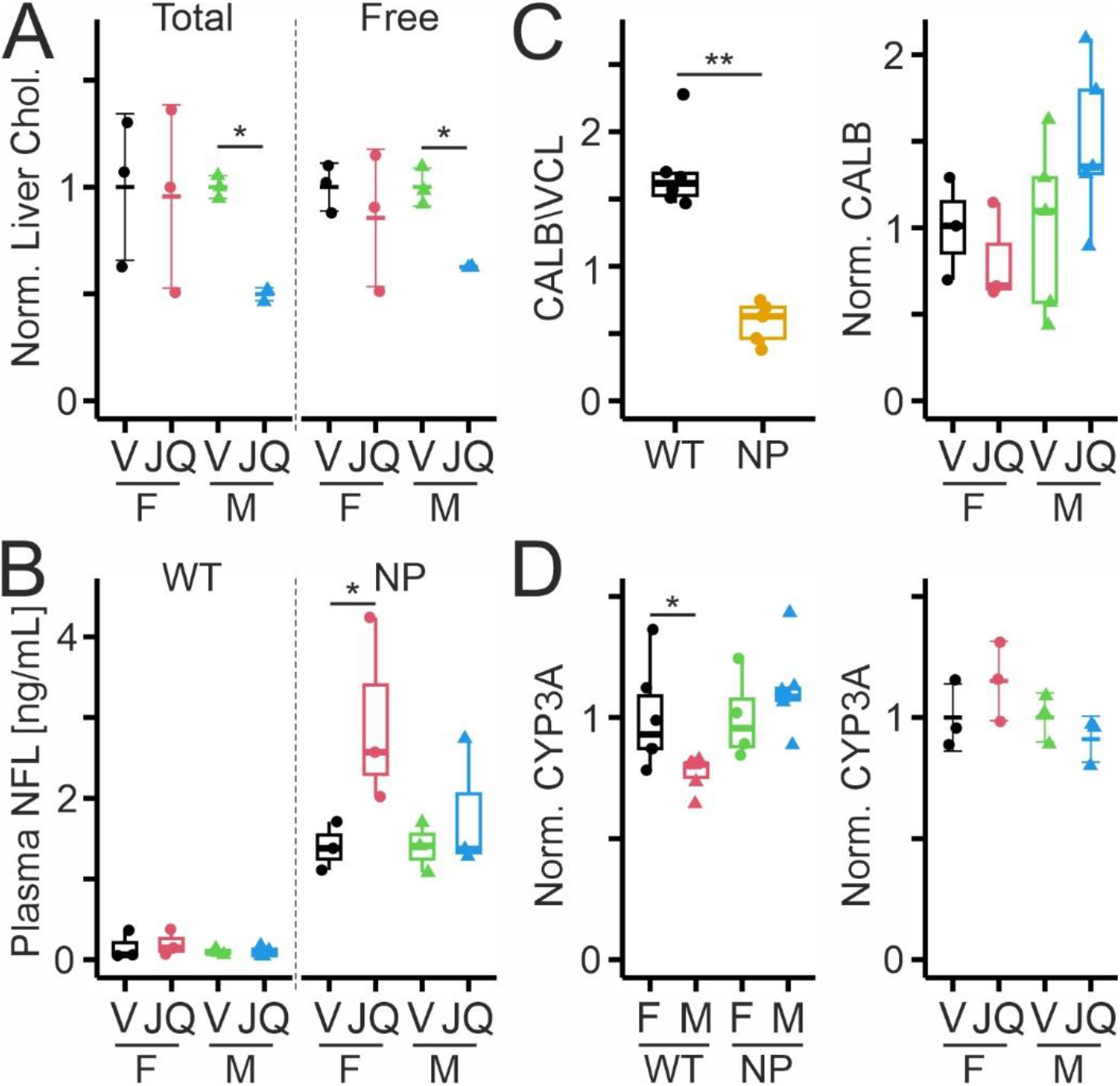
In vivo effects of long-term treatment with JQ1 on molecular markers in selected organs and brain regions of wildtype and mutant mice. A, Normalized levels of total and free cholesterol in liver extracts from female and male mutant mice after treatment for 4 weeks with vehicle (V) and JQ1. Whiskers indicate standard deviations. Asterisks indicate statistically significant differences (*, p < 0.05; Wilcoxon-Mann-Whitney test; n = 3 animals per group). B, Boxplots showing neurofilament light chain (NFL) levels in serum of female (F) and male (M) wildtype (WT) and mutant (NP) mice at 12 weeks of age after 4 weeks of treatment with vehicle or JQ1 Asterisks indicate statistically significant differences (*, p < 0.05; Wilcoxon-Mann-Whitney test; n = 3–4 animals per group). C, Boxplots showing cerebellar protein levels of calbindin (CALB, left) from untreated WT and NP mice at 12 weeks of age normalized to loading control (vinculin, VCL) and (right) from female (F) and male (M) mutant mice after 4 weeks of treatment with JQ1 (red or blue) or vehicle (black or green) normalized first to loading control and then to means of vehicle-treated animals. Asterisks indicate statistically significant differences (**, P < 0.01; Wilcoxon-Mann-Whitney test; genotype: n = 5–6 animals per group; treatment: n = 3–5). D, Boxplots showing protein levels of CYP3A (left) in livers of untreated female and male WT and NP mice at 12 weeks of age and (right) in female and male mutant mice at 12 weeks of age after 4 weeks of treatment with JQ1 or vehicle. Protein levels were normalized first to loading controls and then to means of female mice (left) or of vehicle-treated animals (right). Asterisks indicate statistically significant differences (*, p < 0.05; Wilcoxon-Mann-Whitney test; sex: n = 4–7 animals per group; treatment: n = 3 per group).

## Discussion

Here, we report that BET protein inhibition by JQ1 improved some neurovisceral signs in mice carrying the I1061T variant of NPC1 and that the beneficial changes occurred in male, but not in female animals. Specifically, JQ1 acted on body weight, selected measures of the footprint test, and hepatic cholesterol levels representing core outcome measures for preclinical drug tests in mouse models [52]. In our hands, the footprint and the beam balance test indicated defects in motor coordination of mutant I1061T mice already at 8 weeks of age and showed a positive response to JQ1 treatment. These tests are increasingly used to assess motor control [37, 82] and the efficacy of therapeutic approaches in NPC1 mutant mice [38, 83–86]. In general, previous reports showed variable sensitivity of behavioral tests in different mouse models of NPCD [37, 38, 40, 69, 73, 87, 88]. Notably, our observation that the rotarod test is not sensitive enough to detect impaired motor control in the I1061T model is in agreement with previous reports [31, 37]. Our study underlines the importance to select and validate tests that reliably detect behavioral changes in the mouse strain and colony of interest. Within this context, our application of PCA to measures from different tests appears as highly suitable to identify relevant parameters and to help optimize test designs. As second effect, JQ1 treatment led to an albeit temporary increase in body weight. The temporary nature of this effect is not unusual, it has been observed following treatment with other candidate approaches in mouse models of NPCD [85, 89–92] with drug- and mouse strain-dependent duration of improvement. As third effect, JQ1 reduced hepatic cholesterol accumulation, a visceral hallmark of NPCD that is independent from neurologic symptoms. Previous studies showed similar improvements by hydroxypropyl-beta-cyclodextrin [93, 94], thioperamide [95] and dietary plant stanol [96], whereas other treatments showed no effect [84, 97–99]. At present, it is unclear how JQ1 induces the beneficial effects in male mutant mice. The observed trend to increased calbindin levels in the cerebellum of JQ1-treated mutant mice suggests that JQ1 diminishes loss of Purkinje cells. A similar outcome with improved motor coordination but only moderately increased calbindin levels was observed in NPC1-deficient mice following AAV9-based gene therapy [39] suggesting that measurements of calbindin protein levels are somewhat less sensitive than behavioral tests as readout as they relate only to a subset of neurons. The observed temporary weight gain in male mutant mice may also result from improved motor coordination enabling higher food intake. Previous studies showed that weight loss is induced by neuron-specific NPC1 deficiency in mice [71] and prevented in NPC1-deficient mice by brain-[82] and neuron-specific re-expression of NPC1 [70] indicating a neurologic basis of this key symptom. The decrease of hepatic cholesterol content in male mutant mice supports the hypothesis that JQ1 treatment enhances the protein level of the I1061T variant of NPC1 with residual function and induces a decrease of cholesterol accumulation. However, we did not find robust increases in NPC1 protein in the tissues tested following treatment for 48h or 4 weeks in contrast to our findings in cell culture models [50, 51]. A possible, but speculative explanation is that JQ1 enhanced NPC1 protein levels in vivo only transiently as reflected by the temporary weight gain, at treatment durations when protein levels were not sampled. On the other hand, the observed discrepancy is in line with a previous study showing that treatment with HDAC inhibitors enhanced the protein level of NPC1 in mouse-derived fibroblasts *in vitro* but not in the liver of NPCD mice *in vivo* [100]. Our observation that cMYC protein levels were reduced following long-term treatment both in the visceral organs in brain regions of male mutant mice clearly indicates target engagement of BET proteins. Acting as histone acetylation readers, these proteins affect the expression of numerous genes [101], the observed effects may have been provoked at least in part by NPC1-independent mechanisms. This includes enhancement of autophagy and lysosomal function [102–104], normalisation of cholesterol metabolism [66] as well as anti-inflammatory and neuroprotective effects of BET protein inhibition [105–113]. A remarkable observation is that positive changes were observed in males. This includes weight, footprint test measures, levels of the BET protein target cMYC in different tissues, and hepatic cholesterol. Several studies found evidence for sex-specific effects of bromodomain proteins in pathologic conditions such as epilepsy [114], schizophrenia [115], glioma [116] and cocaine use disorder [117, 118]. However, the underlying mechanisms are not well understood. In the healthy brain, sex-specific expression of genes has been reported recently [119]. BET protein inhibition may impact such transcriptional programs differently. However, the dimorphic effects of JQ1 reported here may be drug-specific, for example due to more rapid elimination of JQ1 by CYP3A in females. Wildtype females showed higher hepatic protein levels than males, but this difference was absent in mutant mice. The sex-dependent effects could also reflect distinct disease progression in males and females. Previous studies noted sex- or gender-specific differences in outcome measures of the disease [69, 120–124], in the impact of modifiers [8] or of drugs [125–127]. The increased serum levels of NFL in female mice occurred only in mutant animals excluding a general neurotoxic effect of the drug in females. It is conceivable that the JQ1-induced increase in NFL levels was caused by enhanced vulnerability of female mutant mice possibly linked to excitotoxicity or neuroinflammation [128–130] compared to males [131]. Alternatively, serum levels of NFL may be affected by enhanced blood coagulation in mutant compared to wildtype mice [132] and they may be influenced by degeneration in the enteric nervous system, but it is unknown whether these changes are sex-dependent.

Our study has several limitations. The I1061T transgenic mice may not be the ideal model due to their specific background and the nature of the transgene. JQ1 is the first generation pan BET protein inhibitor and its pharmacokinetic properties may diminish efficacy. Given the intense interest in these drugs, there are now molecules with higher selectivity, specificity and half-life. Finally, our treatment paradigm, notably the onset and duration of injections, may have been suboptimal to see a full-blown effect of the drug.

## Conclusion

Our proof-of-concept study reveals positive therapeutic effects that call for further in vivo studies exploring BET protein inhibition and its therapeutic potential for NPCD and potentially other lysosomal diseases using other in vivo models, newer inhibitors or degradation enhancers, and different endpoints.

## Declarations

### Ethics approval

The experiments were approved by the Italian Ministry of Health (Rome, Italy; Authorization N° 677/2020-PR) and performed in agreement with the ARRIVE (Animals in Research: Reporting In Vivo Experiments) guidelines, the guidelines of the Italian Ministry of Health (D.L. 26/14) and the European Community Directive 2010/63/EU.

### Consent for publication

NA

### Availability of data and materials

All data generated or analysed during this study are included in this published article and supplementary information. R routines are available on request from the corresponding author (FWP).

## Author’s contributions

Conceptualization: FWP, VP; Data curation: AB, FWP, MP, VP; Formal Analysis: AB, DP, FWP, MP, SC, SL, VP; Funding acquisition: FWP, VP; Investigation: AB, CT, DP, MP, SC, SL, SRF, ST; Methodology: AB, CG, CT, DP, FWP, MP, NCA, SC, SL, SRF, ST, VP; Project administration: FWP, MP, SRF, VP. Resources: AS, CG, FWP, NCA, SRF, VP; Software: FWP; Supervision: FWP, VP; Validation: AB, FWP, MP, SRF, VP; Visualization: FWP; Writing – original draft: FWP; Writing – review & editing: AB, DP, FWP, MP, SC, SRF, ST, VP.

## Supplementary material

Supplementary Figures 1–3, Supplementary Data 1, 2

## Abbreviations

ANOVA: analysis of variance
BET: bromodomain and extra-terminal domain
DMSO: dimethyl sulfoxide
NFL: neurofilament light chain
NP: *Npc1* mutant mice
NPCD: Niemann-Pick type C disease
PC: principal component
PCA: principal component analysis
RT: room temperatureRT
WT: wildtype mice

